# Not all memory declines with age: Neural correlates of implicit associative memory in younger and older adults

**DOI:** 10.64898/2026.10.05.756834

**Authors:** Emily E. Davis, Simon W. Davis, Rosanna K. Olsen, Karen L. Campbell

## Abstract

While explicit associative memory declines with age, implicit associative memory remains preserved. Few studies have examined the neural basis of implicit associative memory in aging, and it remains unclear whether the hippocampus is required as it is more often associated with explicit associative memory. While undergoing functional magnetic resonance imaging, young and old adults completed a speeded object categorization task in which they viewed pictured objects with superimposed words. Implicit memory was tested by presenting intact (original pairs) and rearranged pairs (items previously presented, but not together) from encoding while participants did the same categorization task. Associative priming was defined as faster reaction times (behavioural associative priming) and less activation for intact than rearranged pairs at retrieval (neural associative priming). Whole-brain analyses revealed neural associative priming in occipitotemporal and frontal regions that did not differ by age. A region of interest analysis (using subject-specific masks of medial temporal lobe regions) showed that only older adults had greater posterior hippocampal activation to intact than rearranged items at test, though activation in the hippocampus did not relate to behavioural associative priming. In contrast, greater behavioural associative priming was related to greater neural associative priming in the perirhinal cortex in both age groups. Together, these findings suggest that implicit associative priming remains intact with age, possibly because it relies on additional processes outside the hippocampus such as perirhinal-dependent unitization.

**Significance Statement:** The results from this paper contribute to two debates in the field: first, whether medial temporal lobe regions, especially the hippocampus, support implicit associative memory; and second, whether associative binding is preserved with age. We show that behavioural and neural measures of implicit associative memory are comparable in younger and older adults, and that in older adults, the hippocampus and perirhinal cortex appear to support implicit associative memory. Together, these findings suggest that age-related associative deficits may reflect difficulty with explicit, intentional retrieval rather than a failure to form associations. They also suggest that the hippocampus contributes to relational memory even without conscious awareness. More broadly, this points to implicit measures as a sensitive way to assess memory in aging.

---

A consistent finding in the memory and aging literature is that associative memory shows greater age-related decline than item memory (Old & Naveh-Benjamin, 2008). However, this associative deficit is mainly observed under explicit conditions (e.g., differentiating between intact and rearranged word pairs; Dew et al., 2007). When older adults’ associative memory is tested *implicitly,* deficits are not evident (e.g., they are similarly faster to make a categorization decision for intact than rearranged pairs at retrieval; Davis et al., 2021; Dew & Giovanello, 2010; Kan et al., 2011). This differential performance may be because implicit and explicit associative tasks differ in their dependency on the medial temporal lobes (MTL) and specifically the hippocampus (Slotnick, 2024), a structure affected by aging (Grady & Ryan, 2017). That said, the neural correlates of implicit associative memory have only been examined by a handful of studies (e.g., Gomes et al., 2016; Wang & Giovanello, 2016; Yang et al., 2003). Thus, the current study addresses whether intact implicit associative memory in aging is supported by the hippocampus, or whether it instead relies on regions outside the hippocampus less vulnerable to age-related decline.

Most neuroimaging investigations into implicit memory explore single item priming. These studies show reduced activation to repeated items (i.e., repetition suppression) in occipital, temporal, and frontal regions (Kim, 2019) and the perirhinal cortex (PRC; Wang et al., 2010, 2014), which is largely age-invariant (Bäckman et al., 1997; Lustig & Buckner, 2004; c.f., Ballesteros et al., 2013). In contrast, the neural correlates of implicit *associative* memory are less understood. Historically, the hippocampus was thought to only be required for conscious expressions of associative memory (e.g., Eichenbaum, 2001), as amnesiac patients (with MTL damage) are unimpaired on *implicit* tests of associative memory (Graf & Schacter, 1985; Moscovitch et al., 1986; Verfaelllie et al., 2012; but see Hannula & Greene, 2012). However, this dissociation has been contested (Dew & Cabeza, 2011). Recent work instead ties hippocampal engagement to representational demands rather than memory processes (e.g., Kent et al., 2016), predicting that any binding task, regardless of awareness, should engage the hippocampus. Indeed, both functional neuroimaging (Degonda et al., 2005; Gomes et al., 2016; Henke et al., 2003) and lesion studies (Carlesimo et al., 2005; Yang et al., 2003) point to hippocampal involvement in implicit associative memory.

Given that hippocampal structure and function decline with age and affect explicit associative memory (Becker et al., 2015; Carr et al., 2017; Shing et al., 2011), it is surprising that behavioural measures of associative priming remains intact (E. Davis et al., 2021; Dew & Giovanello, 2010). This could be because explicit tests place additional demands on frontally-mediated control mechanisms that also decline with age (e.g., Brehmer et al., 2020; Guardia et al., 2023). By this account, hippocampal binding may be preserved in aging (Campbell & Davis, 2024). Consistent with this, the one neuroimaging study that investigated implicit associative memory in aging found that the posterior hippocampus showed increased activation for intact compared to rearranged pairs of words at retrieval in both younger and older adults (Wang & Giovanello, 2016). However, their paradigm differed from traditional priming tasks in ways that limit what can be inferred about associative priming per se. The word pairs were embedded in sentences; thus, any priming benefit may reflect reinstatement of a shared semantic context rather than a direct link between the two items. Sentences also provide the kind of semantic scaffolding known to minimize age differences in associative memory (Naveh-Benjamin et al., 2007). To make claims about associative priming, as opposed to conceptual priming, the task should require the generation of a link between two otherwise unrelated concepts.

Here, we tested whether intact implicit associative memory in aging is supported by the hippocampus using a speeded object categorization task (E. Davis et al., 2021; Gomes et al., 2016). Whole-brain univariate analyses identified regions sensitive to neural associative priming (rearranged > intact pairs; Grill-Spector et al., 2006). These were complemented by a region-of-interest (ROI) analysis targeting the hippocampus, using subject-specific ROIs derived from an automated segmentation of each participants’ T1-weighted images. We additionally examined the parahippocampal cortex and perirhinal cortex, which have been implicated in fMRI studies on associative (Yang et al., 2008) and item priming (Voss et al., 2009; Wang et al., 2014; Wang & Giovanello, 2016), respectively.

## Method

### Participants

Participants were recruited by poster and newspaper advertisement. Data was collected from 30 young adults (18-29 years) and 30 old adults (60-80 years). Data were excluded for the following reasons: withdrew from the study (YA n = 1), scored less than 23 on the Montreal Cognitive Assessment (MoCA; OA n = 2; Carson et al., 2018; Nasreddine et al., 2005), and excessive scanner motion (YA n = 1; OA n = 3; described below). These exclusions resulted in a final sample of 25 older adults and 28 younger adults.

Older adults were more highly educated, *t* (51) = 4.25, *p* < .001, and had higher vocabulary scores, *t* (50) = 6.66, *p* < .001, than young adults. See Table 1 for demographic information.

**Table 1.** Demographics.

|  | **Young** | **Old** |
| --- | --- | --- |
| Age | 21.20 (2.76) | 70.40 (5.83) |
| Male/Female | 7/21 | 12/12 |
| MoCA |  | 28.20 (2.27) |
| Shipley Vocabulary Test | 30.0 (3.54) | 36.30 (3.23) |
| Education years | 15.80 (1.92) | 18.60 (2.77) |

*Note.* Sex (at birth) was missing for one older adult. Vocabulary score was missing for one young adult.

### Stimuli

Stimuli were from Davis et al. (2024). One-hundred ninety-six images of real-life, everyday objects (300 × 300 px) were selected from Hovhannisyan et al. (2021). Another 196 unique object-words were generated from other objects in the database that were not being used as pictures (e.g., a picture of a cardinal would generate the object word “bird”). These pictures and words were evenly split into living and non-living items and then randomly paired with some restrictions: Living pictures were paired with non-living words (and vice versa), and 50% of the pairs needed to generate a yes response to the object categorization question (i.e., could the object(s) fit inside a common desk drawer). To ensure that participants always assessed both the picture and the word when asked to attend to both, the pairings were not always congruent (i.e., the picture did not always fit if the object-word did). This resulted in 16 incongruent and 28 congruent pairs in each list.

Once the images were paired, they were split into four lists of 44. The remaining 24 pairs were used for fillers at the beginning of the tasks (three per task) and practice trials (n = 8). For the memory test, the pairs were split evenly into 22 intact and 22 rearranged pairs. For rearranged items, the objects were re-paired such that responses to the object-categorization question remained the same from encoding to test to ensure that any changes in reaction time were a result of changes to the stimulus pairs and not to the stimulus-response binding (Gomes & Mayes, 2020). The lists were counterbalanced across participants such that each participant saw two of four possible lists (88 total trials) that were shown in separate runs. Stimuli within these lists were counterbalanced in both the intact and rearranged conditions. The number of incongruent and congruent pairs were evenly split between the intact and rearranged conditions. Word length and frequency were matched across the four lists and pair type conditions.

### Procedure

The experiment was approved and carried out in line with the requirements instituted by the Office of Research Ethics at Brock University.

#### Scanner: Day 1

The implicit associative memory paradigm was adapted from Davis et al. (2021, 2024; originally Dew & Giovanello, 2010) and presented using E-prime 3 (Psychology Software Tools, Pittsburgh, PA, USA). Prior to entering the scanner, participants practiced eight trials of the object categorization task and had the opportunity to ask questions or repeat the task if necessary. Once participants were in the scanner, they completed the same practice as earlier once more. Then, two encoding-test blocks of an object categorization task presented in four separate scanner runs. In both blocks, participants first completed an incidental encoding phase, in which they were shown pictures of everyday objects with superimposed yellow object-words (presented for 2,000 ms; see Fig. 1), separated by a 500 ms ISI and a jittered fixation cross (*M* = 3075 ms). Participants were instructed to look at the pictured object and word to answer whether the objects could fit together inside of a common desk drawer. Speed and accuracy were emphasized. All pairs were presented twice at encoding such that they were presented once in random order before they were presented the second time. No pairs were shown twice in sequence (i.e., the first item in the second presentation was never the last item in the first presentation). Following encoding, participants completed an implicit associative memory test. In the test block, participants viewed the same objects and words from the encoding block, but unbeknownst to the participants, some of the pairs at test were intact (i.e., the same pairing from encoding) and some were rearranged (i.e., pictures and words previously presented, but not together). The behavioural outcome of interest for these tasks was reaction time (RT). Participants also completed two other encoding-test blocks with different stimuli and encoding instructions (data not reported here).

**Figure 1.**
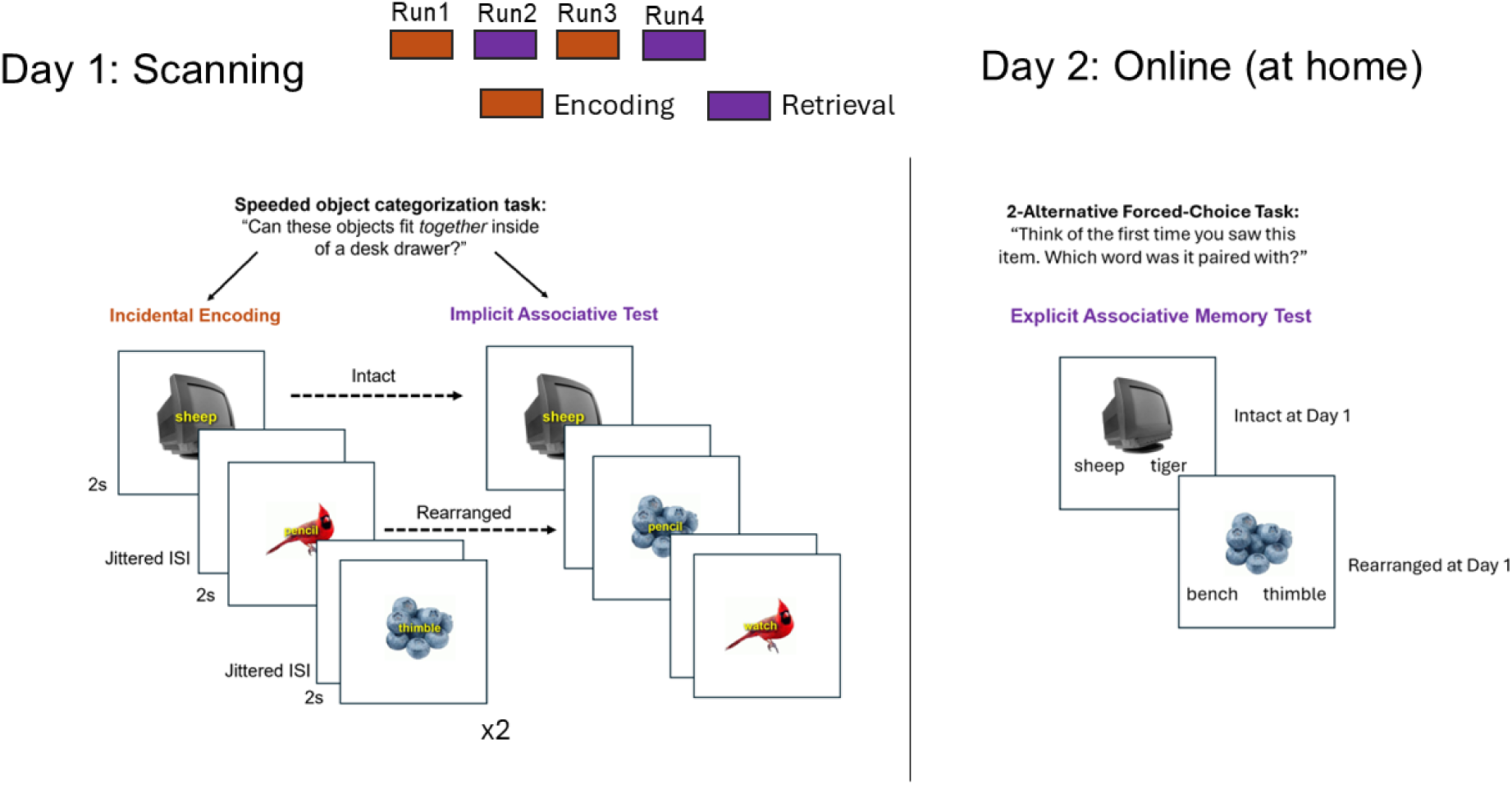
Experimental Procedure. *Note.* Participants completed a two-part study. On Day 1, they were scanned while they did two encoding-test blocks of the implicit associative memory task. On Day 2, they completed a 2-Alternative Forced-Choice Task at-home online. They saw all the pictures from Day 1 and had to decide which word it was paired with. The word choices were always from the original pairing at encoding and familiar word seen at encoding.

Outside of the scanner, participants completed the MoCA (Nasreddine et al., 2005), Shipley Vocabulary Test (Shipley & Burlingame, 1941), and a demographics form.

#### At-home Online: Day 2

To test whether older adults showed an age-related deficit in explicit associative memory, we probed memory for the implicitly learned pairs on Day 1 in an online task the day after scanning. The Day 2 task was programmed with jsPsych (version 7.3.2; de Leeuw, 2015) and hosted on an Amazon Web Server. Participants were not told that the task was a memory test; only that they would continue doing a similar experiment. Participants were sent a link to the online experiment the following morning and were asked to complete the task before the end of the calendar day. In the task, participants viewed the same pictured-objects from the day before with two words underneath in black text. They were asked to select which word was paired with the picture the previous day. One of the words was the word that was paired with the object at encoding, and the other word was another word from the task but not presented with that object on Day 1. Thus, both words were equally familiar to the participant. We tested all objects from Day 1, which means that some of the items were *intact* during the implicit associative test and some were *rearranged.* The correct answer was the word that it was paired with during the *encoding* phase, and the rearranged pairs themselves were not tested. The correct answer was presented on the left side of the screen in half of the trials (see Fig. 1).

### MRI Acquisition

Participants were scanned at the Toronto Neuroimaging Center. A high-resolution T1-weighted anatomical image was acquired using a magnetization-prepared rapid gradient-echo (MPRAGE) sequence (TR = 2500 ms, TE = 4.37 ms, TI = 1100 ms, flip angle = 7°, FOV = 256 mm, 192 sagittal slices, 1.0 × 1.0 × 1.0 mm³ isotropic voxels, GRAPPA acceleration factor = 3). Functional images were acquired using a T2*-weighted echo-planar imaging (EPI) sequence with simultaneous multi-slice acceleration (TR = 2000 ms, TE = 30 ms, flip angle = 70°, FOV = 220 mm, 69 interleaved axial-oblique slices angled −20° relative to the AC–PC line, 2.0 × 2.0 × 2.0 mm³ isotropic voxels, SMS/multiband factor = 3). To correct for magnetic field inhomogeneities, two dual-echo gradient-echo field maps were acquired (TR = 646 ms, TE1 = 4.92 ms, TE2 = 7.38 ms, flip angle = 45°, same FOV, slice prescription, and resolution as the functional data: 220 mm FOV, 69 slices, 2.0 × 2.0 × 2.0 mm³ voxels).

### MRI Preprocessing

Data were converted to BIDS using dcm2bids (3.1.1; Bore et al., 2023). At this time, fieldmap scans were assigned to the two or three scans that came before and after the fieldmap was acquired. Anatomical and functional quality control was done with MRIQC (Esteban et al., 2017). Output from MRIQC was manually inspected to check for artifacts and evaluate participant motion. Participants were identified as outliers if they had FD > .50 mm for > 50% of the scan (1 YA and 3 OAs removed, as mentioned above). Older adults (*M* = 0.22, *SD* = 0.10) had larger mean framewise displacement than younger adults (*M* = 0.14, *SD* = 0.05), *t* (42.43) = 4.09, *p* < .001. Data were then submitted to fMRIPrep (Esteban et al., 2019; see boilerplate below) and then smoothed with a 8 mm kernel in line with Gomes et al. (2016).

### fMRIPrep Preprocessing

Data were preprocessed with *fMRIPrep* 25.2.4 (Esteban et al. (2019); Esteban et al. (2018); RRID:SCR_016216), which is based on *Nipype* 1.10.0 (K. Gorgolewski et al. (2011); K. J. Gorgolewski et al. (2018); RRID:SCR_002502).

### Preprocessing of B_0_ inhomogeneity mappings

For each of the two fieldmaps, a *B_0_* nonuniformity map was estimated from the phase-drift maps measure with two consecutive GRE (gradient-recalled echo) acquisitions. The corresponding phase-map(s) were phase-unwrapped with prelude (FSL None).

### Anatomical data preprocessing

The T1w image was corrected for intensity non-uniformity (INU) with N4BiasFieldCorrection (Tustison et al. 2010), distributed with ANTs 2.6.2 (Avants et al. 2008, RRID:SCR_004757), and used as T1w-reference throughout the workflow. The T1w-reference was then skull-stripped with a *Nipype* implementation of the antsBrainExtraction.sh workflow (from ANTs), using OASIS30ANTs as target template. Brain tissue segmentation of cerebrospinal fluid (CSF), white-matter (WM) and gray-matter (GM) was performed on the brain-extracted T1w using fast (FSL (version unknown), RRID:SCR_002823, Zhang, Brady, and Smith 2001). Volume-based spatial normalization to one standard space (MNI152NLin2009cAsym) was performed through nonlinear registration with antsRegistration (ANTs 2.6.2), using brain-extracted versions of both T1w reference and the T1w template. The following template was selected for spatial normalization and accessed with *TemplateFlow* (25.0.4, Ciric et al. 2022): *ICBM 152 Nonlinear Asymmetrical template version 2009c* [Fonov et al. (2009), RRID:SCR_008796; TemplateFlow ID: MNI152NLin2009cAsym].

### Functional data preprocessing

First, a reference volume was generated, using a custom methodology of *fMRIPrep*, for use in head motion correction. Head-motion parameters with respect to the BOLD reference (transformation matrices, and six corresponding rotation and translation parameters) are estimated before any spatiotemporal filtering using mcflirt (FSL , Jenkinson et al. 2002). The estimated *fieldmap* was then aligned with rigid-registration to the target EPI (echo-planar imaging) reference run. The field coefficients were mapped on to the reference EPI using the transform. The BOLD reference was then co-registered to the T1w reference using mri_coreg (FreeSurfer) followed by flirt (FSL, Jenkinson and Smith 2001) with the boundary-based registration (Greve and Fischl 2009) cost-function. Co-registration was configured with six degrees of freedom. BOLD runs were slice-time corrected to 0.948s (0.5 of slice acquisition range 0s-1.9s) using 3dTshift from AFNI (Cox and Hyde 1997, RRID:SCR_005927). Several confounding time-series were calculated based on the *preprocessed BOLD*: framewise displacement (FD), DVARS and three region-wise global signals. FD was computed using two formulations following Power (absolute sum of relative motions, Power et al. (2014) and Jenkinson (relative root mean square displacement between affines, Jenkinson et al. (2002). FD and DVARS are calculated for each functional run, both using their implementations in *Nipype* (following the definitions by Power et al. 2014). The three global signals are extracted within the CSF, the WM, and the whole-brain masks.

### Experimental Design and Statistical Analysis

#### Behavioural Analysis

Behavioural data were processed in R with the rprime (Mahr, 2025) and tidyverse (Wickham et al., 2019) libraries. Object categorization accuracy and reaction times (RT) were the behavioural outcome measures. Before averaging participant responses, RT data were trimmed such that anticipatory responses less than 200 ms were removed, then RTs greater than 2.5 SD from an individual’s mean for each condition were removed. Only accurate responses were used to calculate mean RTs.

To ensure that participants learned the items, we first checked that participants responded faster to pairs with repeated presentation. To do this, we entered RTs from the first- and second-time participants saw the pairs at encoding (Time 1 & 2) and the *intact pairs* from the implicit test (Time 3) into a 2 (Age Group: young, old) x 3 (Time: 1, 2, 3) ANOVA. To test the primary question of whether there were age differences in behavioural *associative* priming (rearranged > intact RTs), the RT data from the implicit associative memory test were submitted to a 2 (Age Group: young, old) x 2 (Pair Type at Day 1 Test: intact, rearranged) mixed ANOVA.

To explore individual differences in the relationship between brain and behaviour, we calculated behavioural priming scores, defined as faster responding to intact than rearranged pairs at test. Instead of using raw difference scores, which are known to be more variable than the individual means that go into them (Draheim et al., 2019) and to account for age differences in baseline RT, we regressed mean RTs for intact pairs onto rearranged pairs across participants for each block type and used the residual as an implicit associative priming score for that individual (E. E. Davis et al., 2024, 2025; Healey et al., 2014). Higher positive values indicate greater binding (i.e., faster responding to intact than rearranged pairs). These behavioural measures were used as dependent variables in linear regressions with Age Group and ROI-level parameter estimates (betas from the rearranged > intact contrast) as predictors

#### MRI Analysis

Model specification for the implicit associative memory task was done in line with Gomes et al., (2016) whose design we closely followed. Effects were estimated using a General Linear Model (GLM) in which the BOLD response to each event-type was modeled by convolving a delta function at stimulus onset with a canonical hemodynamic response function. Only trials with accurate responses to the object categorization task were included. A GLM was created for each retrieval run, including regressors representing the events (intact, rearranged), as well as the parametric modulation of the RTs for each event using a polynomial expansion of the first order. Additional covariates of no interest included six regressors representing the movement parameters estimated during realignment, session effects, and a global mean.

##### Whole Brain Analysis

For the whole-brain voxel-wise analyses, data were thresholded at an FDR corrected cluster-level of p < 0.05 (with a cluster-forming threshold of p < 0.001). Localization of significant peaks was done using standard anatomical atlases (AAL). Priming effects are generally associated with neural repetition suppression (i.e., decreased activation to repeated pairs; Grill-Spector et al., 2006). As such, to measure neural associative priming, our contrast of interest was greater activation to rearranged pairs compared to intact pairs at test (Gomes et al., 2016; Yang et al., 2008). However, it is worth noting that Wang & Giovanello (2016) found that there was greater activation for intact compared to rearranged trials in the hippocampus, so we test this contrast as well.

##### ROI Analysis

To explore the specific contribution of the hippocampus to associative priming, we also performed a ROI analysis with subject-specific MTL subregion masks, which included the hippocampus and MTL cortex. To create the subject-specific masks, T1-weighted anatomical scans were submitted to the Automatic Segmentation of Hippocampal Subfields (ASHS) software using the ASHS-PMC-T1 atlas (Xie et al., 2016). This software automatically segments participants’ T1 scan into the anterior and posterior hippocampi as well as other medial temporal lobe regions, including the parahippocampal cortex (PHC) and perirhinal cortex (PRC). See Figure 3a. Binary ROI masks were created using Convert3D in ITK-SNAP (Yushkevich et al., 2006; www.itksnap.org), then normalized to MNI space using the warping coordinates from fMRIPrep’s T1 → MNI space transformation (.image_xfm.h5) with *AntsApplyTransforms* (Avants et al., 2009; v.2.6.0) in Neurodesk (Renton et al., 2024). These masks were down sampled from 1 mm to 2mm (ANTs: *ResampleImage*) to match the resolution of the functional data. These subject-specific masks were used to extract mean activation levels from the functional data for the rearranged > intact contrast. All segmentations were manually inspected in both native and MNI space to ensure the segmentations fell within anatomically plausible locations.

**Figure 2.**
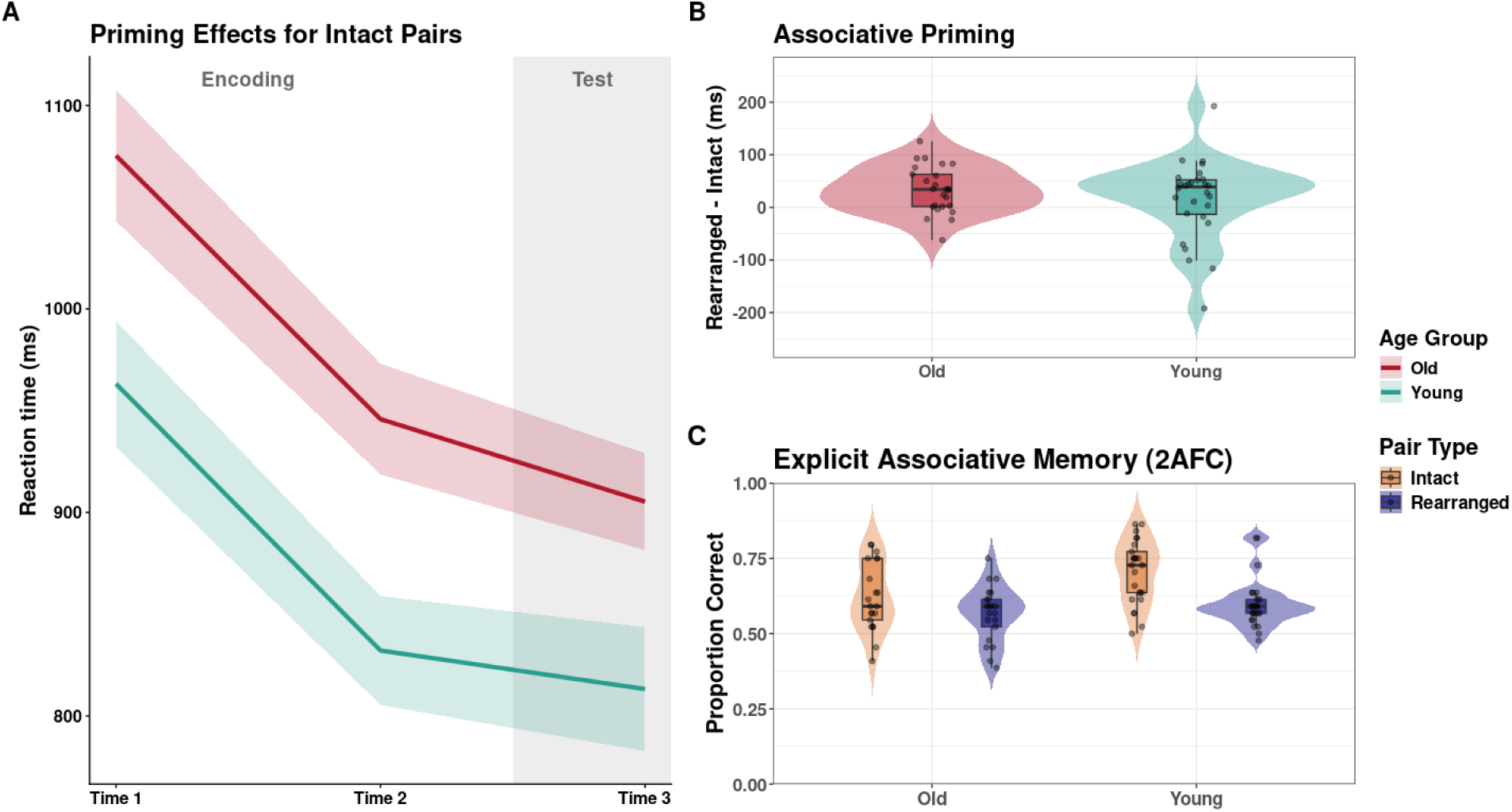
Behavioural Results for Implicit and Explicit Measures of Memory. *Note*. A) Reaction times (RTs) across repetitions of intact pairs. Time 1 and Time 2 reflect the first and second presentations of each pair during encoding. Time 3 reflects the third presentation of the intact pair during the implicit associative memory test. B) Associative priming scores calculated as rearranged RT minus intact RT. Larger values indicate greater associative priming. C) Explicit associative memory performance (forced-choice) assessed on Day 2. Pair type reflects whether each pair appeared intact or rearranged during the implicit associative memory test on Day 1.

**Figure 3.**
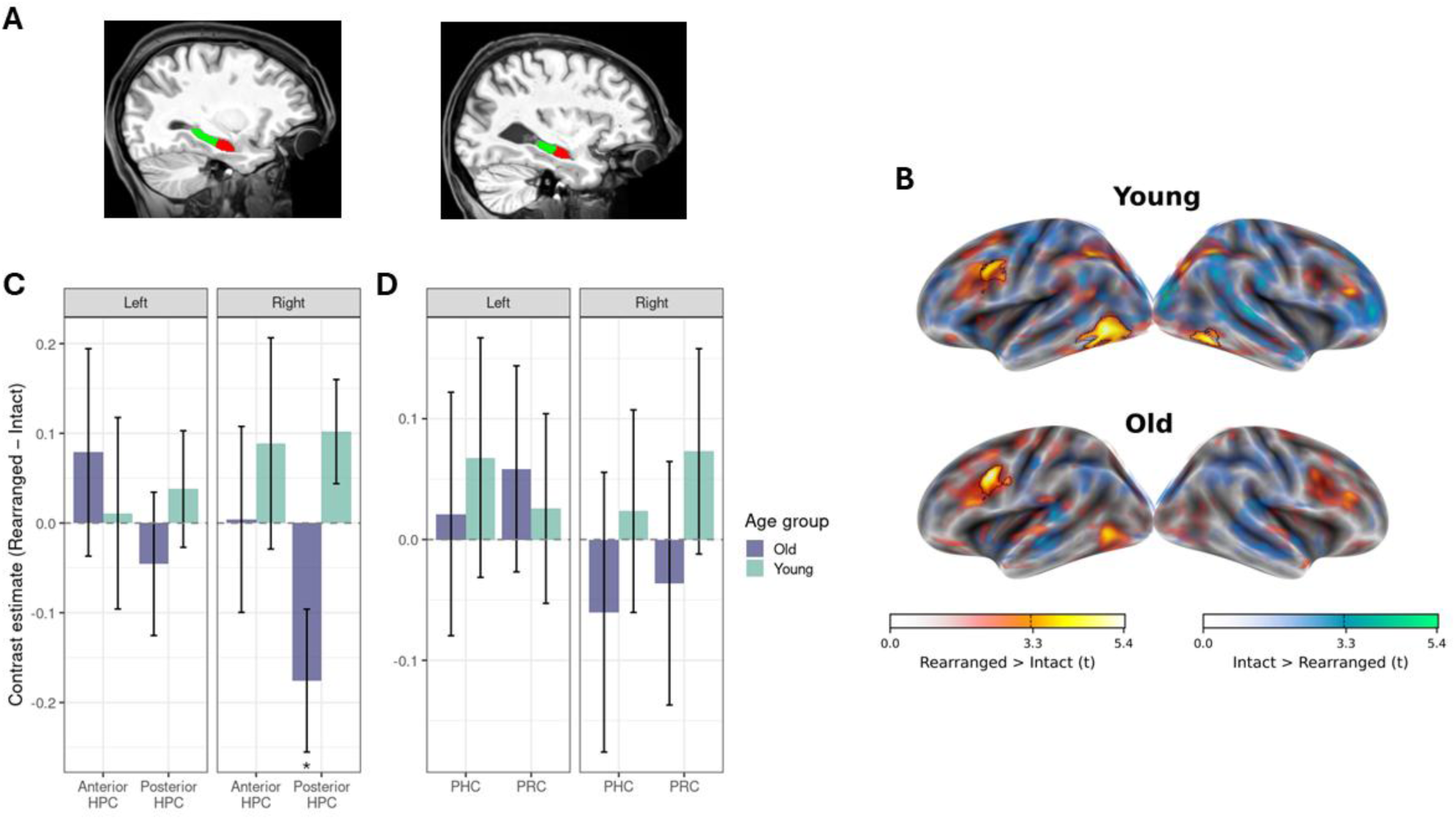
Hippocampal and Medial Temporal Lobe Responses to Rearranged Versus Intact Pairs in Younger and Older Adults. *Note.* A) Subject-specific hippocampal masks for a representative younger (left) and older (right) adult. Red = anterior, green = posterior. B) Group-level *t*-statistic maps for the Rearranged > Intact and Intact > Rearranged contrasts. Maps are displayed with transparent thresholding, which retains subthreshold context rather than masking it (Taylor et al., 2026). Each surface was rendered with an opacity that scaled with its statistic, while subthreshold faces faded quadratically toward zero opacity. Regions surviving cluster-level correction are outlined in black. Note that no regions showed greater activation for younger or older adults, but age groups are plotted separately for illustrative purposes. Warm colours = Rearranged > Intact contrast; cool colours = Intact > Rearranged contrast. C) ROI analysis results. Error bars represent SE of the mean. Positive values represent greater activation for Rearranged than Intact trials. Mean contrast estimates for the hippocampus (HPC) separated by hemisphere and anterior and posterior axis. D) Mean contrast estimates for the parahippocampus (PHC) and perirhinal cortex (PRC). * *p* < .05.

## Results

### Behaviour

#### Accuracy

We first evaluated whether participants were making the correct categorization responses at both encoding and test in the implicit task (i.e., would the objects fit in a desk drawer together or not). Accuracy for both older and younger adults was greater than 85%. For the encoding phase, accuracy did not differ between older (*M* = .90, *SD* = .05) and younger adults (*M* = .89, *SD* = .06), *t* (51) = .88, *p* = .379. For the test phase, accuracy was submitted to a 2 (Age group: young, old) x 2 (Pair Type: intact, rearranged) ANOVA. This analysis also indicated that there were no differences in accuracy across Age Group, *F* (1,51) = 0.07, *p* = .800, pair type *F* (1,51) = 1.30, *p* = .260, nor did these factors interact, *F* (1,51) = 0.25, *p* = .619 (Old: *Mintact* = .91, *SD* = .07; *Mrearranged =* .92, *SD* = .04; Young: *Mintact* = .91, *SD* = .05; *Mrearranged =* .92, *SD* = .04).

#### Implicit Associative Priming

To confirm that participants showed behavioural evidence of priming for intact pairs from encoding to test, we entered RTs from the first and second time participants saw the pairs at encoding (Time 1 & 2) and the *intact pairs* from the implicit test (Time 3) into a 2 (Age Group: young, old) x 3 (Time: 1, 2, 3) ANOVA. This yielded a significant effect of Age Group, *F* (1,51) = 7.33, *p* = .009, such that older adults responded more slowly than young adults. There was a main effect of Time, *F* (2,102) = 137.12, *p* < .001, but no Age Group x Time interaction. Pairwise comparisons (Bonferroni corrected) support that this main effect of Time was driven by faster responses from Time 1 to Time 2, *t* (102) = 12.68, *p* < .001, and from Time 2 to Time 3, *t* (102) = 2.90, *p* = .01 in both age groups. See Figure 2a and Table 2 for RT means.

**Table 2.**
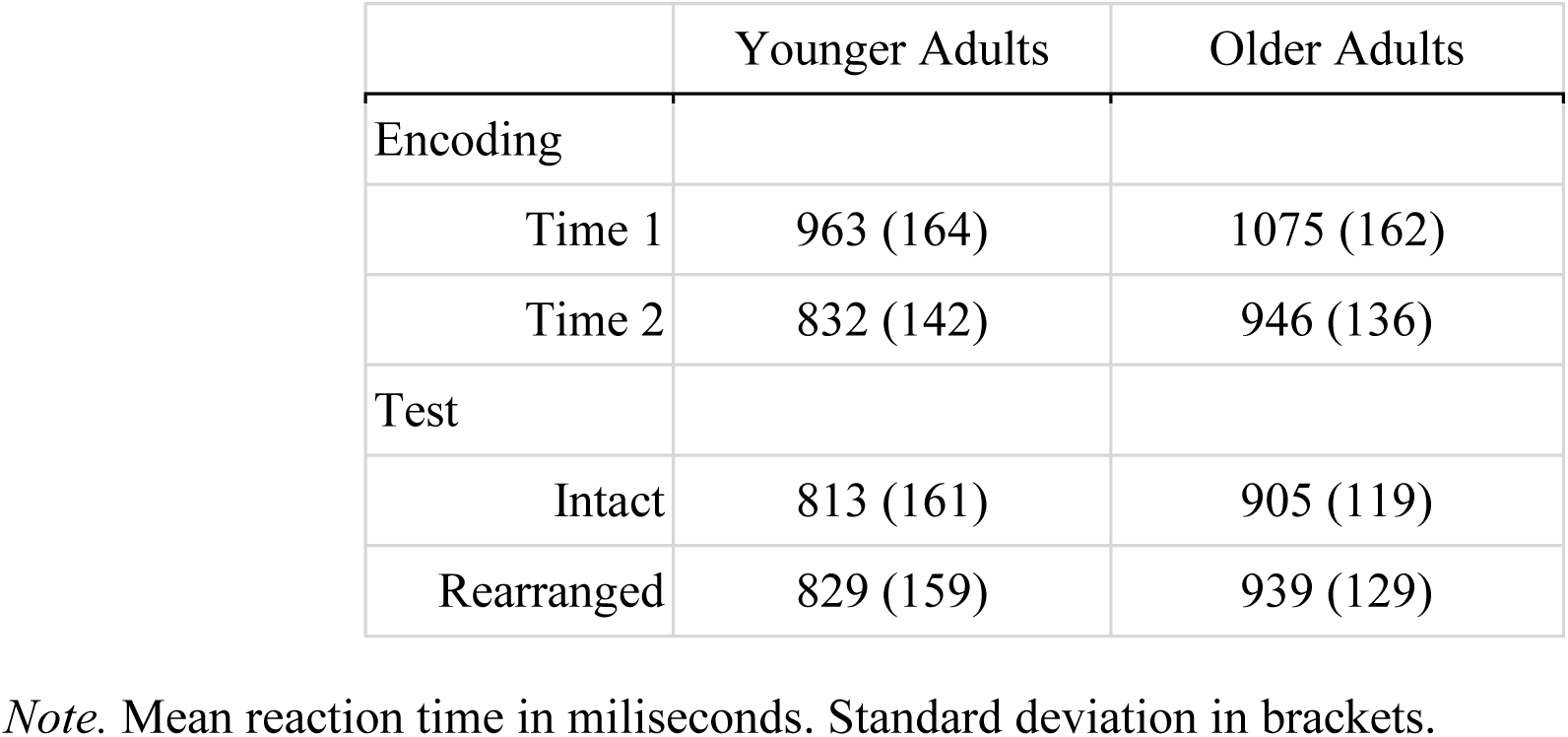
Reaction Times for the Implicit Associative Memory Task.

The primary behavioural effect of interest was the replication of past studies that have shown significant *associative* priming (faster responding to intact than rearranged pairs at retrieval) in both young and older adults (Dew & Giovanello, 2011; Kan et al., 2011; but see Davis et al., 2021 who showed this effect only in older adults). To test this, the RT data were entered into the 2 (Age group: young, old) x 2 (Pair Type: intact, rearranged) ANOVA. This yielded a main effect of Age Group, *F* (1,51) = 6.80, *p* = .012, such that older adults responded more slowly than young adults. There was a main effect of Pair Type, *F* (1,51) = 7.93, *p* = .007, such that participants were faster to respond to intact than rearranged trials. Critically, there was no Age Group x Pair Type interaction, *F* (1,51) = 1.07, *p* = .307, suggesting that associative priming did not differ with age (Fig. 2b). These results support past work showing intact implicit associative memory in aging (E. Davis et al., 2021; Dew & Giovanello, 2010).

#### Next Day Explicit Memory

Three older and three younger adults did not complete Day 2. One additional older adult participant was excluded from the analysis for experimenter error (old n = 21; young n = 25). To test for age differences in explicit memory, accuracy from the 2-alternative forced choice task completed on day two was entered into a 2 (Age Group: young, old) x 2 (Pair Type at Day 1 Test: intact, rearranged) ANOVA. This yielded a significant effect of Age Group, *F* (1, 44) = 6.29, *p* = .016, such that older adults (*M* = .59, *SD* = .12) had worse accuracy than younger adults (*M* = .65, *SD* = .12). There was also a significant effect of Pair Type, *F* (1, 44) = 22.25, *p* < .001, such that there was worse accuracy to the pairs that were rearranged during the implicit memory test (*M* = .58, SD = .08) than the pairs that remained intact (M = .67, SD = .08), as expected based on existing work on the role of interference in memory (MacLeod, 2024 for review). There was no Age Group x Pair Type interaction, *F* (1, 44) = 2.16, *p* < .148 (see Fig. 2c). Importantly, when both age groups’ mean accuracy was tested against chance in a one-sample t-test (*mu* = .50), both age groups performed significantly above chance, *p’s* < .001, indicating that both groups learned the pairs from the day prior. These results support past literature on age-related deficits in explicit associative memory.

### Functional MRI Results

#### Whole-brain Univariate Analysis

Across both age groups, there was evidence of significant neural associative priming (i.e., intact < rearranged) in several regions. Specifically, there was greater activation in clusters with peak activation in the left precentral gyrus (cluster extends into the left inferior pars triangularis and pars opercularis), left inferior temporal gyrus (cluster extends into the left inferior occipital, left fusiform, and middle temporal lobe), and right middle frontal gyrus (see Table 3 and Figure 3). To explore whether hippocampal activation could be detected at a more liberal threshold, the cluster threshold is dropped to k = 20. There was additional activation in the right middle occipital lobe, right frontal inferior frontal gyrus (pars opercularis) and right inferior temporal. No clusters showed greater activation for younger or older adults, even uncorrected, *ps* > .10, but activation maps are shown for older and younger adults separately in Figure 3b for illustrative purposes.

**Table 3.** Whole Brain Univariate Results.

| **Contrast** | **Region** | **p~FDR**<br>**(cluster)** | **k** | **t** | **x** | **y** | **z** |
| --- | --- | --- | --- | --- | --- | --- | --- |
| Rearranged > Intact | L Precentral | 0.000 | 829 | 6.82 | -36 | 8 | 32 |
|  | L Inferior Frontal |  |  | 3.95 | -44 | 24 | 22 |
|  | L Inferior Temporal | 0.000 | 645 | 6.37 | -52 | -62 | -6 |
|  | L Inferior Temporal |  |  | 3.36 | -48 | -58 | -22 |
|  | R Middle Frontal | 0.041 | 168 | 4.31 | 40 | 32 | 20 |
| Intact > Rearranged | R Middle Temporal | 0.000 | 301 | 4.41 | 66 | -30 | 0 |
|  | R Inferior Temporal |  |  | 4.32 | 55 | -22 | -18 |
|  | R Middle Temporal |  |  | 4.24 | 56 | -32 | -4 |
|  | R Cuneus | 0.000 | 284 | 4.34 | 8 | -74 | 26 |
|  | R Cuneus |  |  | 4.19 | 4 | -84 | 22 |
|  | L Cuneus |  |  | 3.89 | -6 | -86 | 26 |

Note. k = cluster extent in voxels; coordinates in MNI space; p-values FDR-corrected at cluster level. Results are across both age groups.

For the intact > rearranged contrast, there was significant activation in the right middle temporal lobe (cluster extends into the right inferior and superior temporal lobe) and the right cuneus (see Table 3 and Figure 3b). To determine if hippocampal activation was evident at a liberal threshold, the cluster threshold was dropped again to k = 20. At this threshold, additional clusters were evident in the right superior medial frontal lobe, the left middle temporal lobe, left supramarginal gyrus, right angular gyrus, and right anterior cingulum. No clusters showed greater activation for younger or older adults (uncorrected, *ps* > .10). Together, there was no evidence that the hippocampus differentiated between intact and rearranged pairs in the whole-brain analyses.

#### ROI Analysis

The hippocampus did not emerge in either whole-brain contrast at the neither the chosen statistical threshold nor at a lower threshold. To further probe whether this structure was engaged in associative priming, we conducted a ROI analysis. Because Wang and Giovanello (2016) reported age differences along the anterior–posterior hippocampal axis, we extracted mean beta estimates for the rearranged > intact contrast within subject-specific anterior and posterior hippocampal ROIs in each hemisphere. A one-way ANOVA with age group as the independent variable was run for each ROI. The only analysis that yielded a significant result was for the right posterior hippocampus, *F* (1,52) = 8.10, *p* = .006, η²p = .135, which was driven by older adults showing greater activation for intact pairs than rearranged pairs in the right posterior hippocampus (*M* = -.18, *SD* = .41), compared to young adults who showed greater activation for rearranged pairs compared to intact (*M* = .10, *SD* = .31). This difference was significantly different from zero in the older adults, *t* (25) = 2.21, *p* = .037, but not the younger adults, *t* (27) = 1.76, *p* = .090. No other hippocampal region approached significance (all *F*s < 0.67, all *p*s > .42, all η²p ≤ .013; see Fig. 4a).

**Figure 4.**
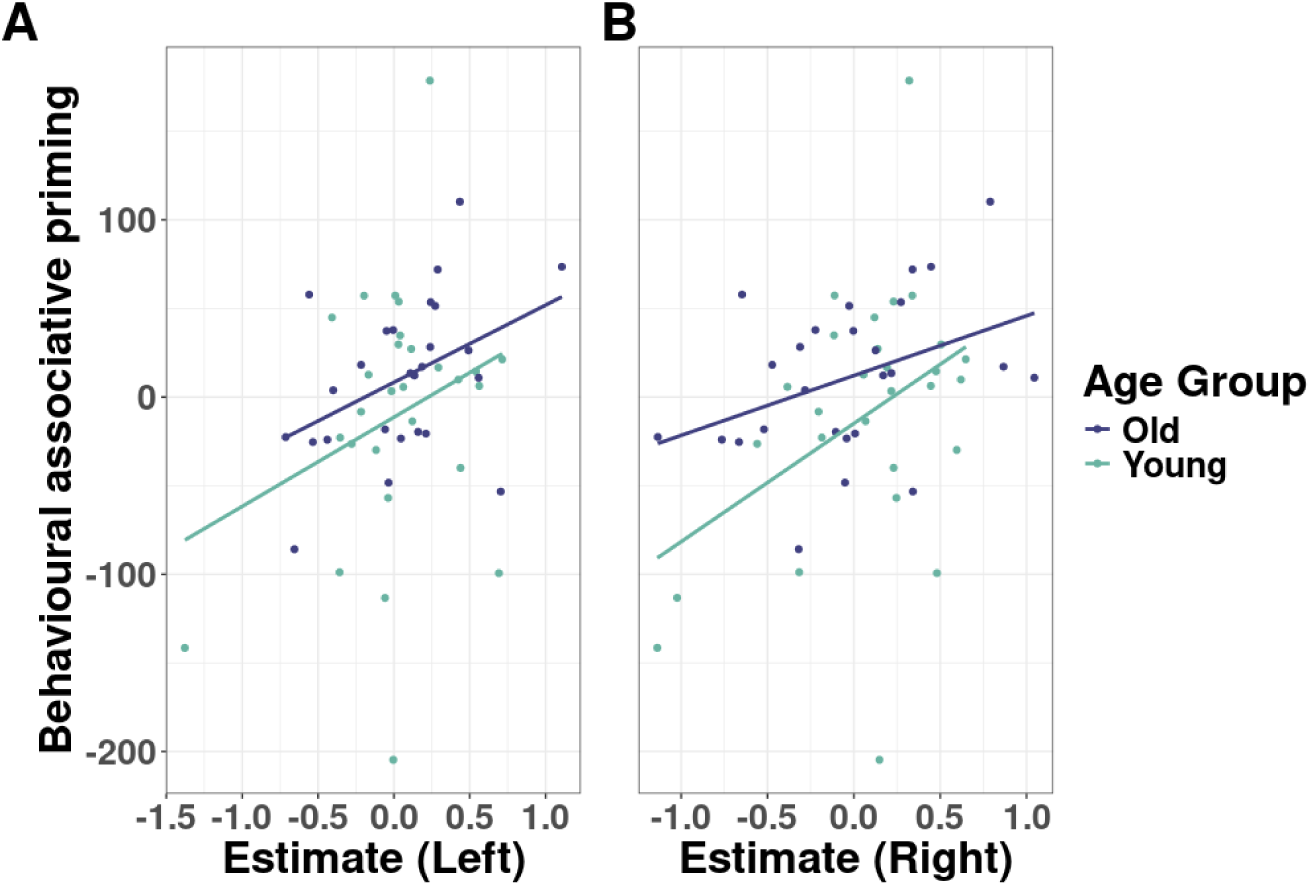
Neural associative priming in the PRC relates to behavioural associative priming measures. *Note.* The relationship between behavioural associative priming (rearranged – intact; larger values indicate greater priming) and neural associative priming in the left (A) and right (B) PRC (rearranged > intact contrast; larger values indicate greater neural associative priming).

ROIs for the PHC and PRC were also tested due to past work implicating these regions in associative priming (Yang et al., 2008) and PRC in item priming (Wang et al., 2014), respectively. However, none of these analyses reached significance (all *F*s < 0.69, all *p*s > .40, all η²p ≤ .013; see Fig. 4b).

To determine whether neural associative priming related to behavioural associative priming, we ran separate regressions predicting behavioural priming from Age Group and the parameter estimates from each MTL ROI from the neural priming contrast (rearranged > intact). The only relationship that was significant was in the left and right PRC (left: *b* **=** 46.95, *p* = .019; right: *b* **=** 16.58, *p* = .004; see Supplemental Table 1 for all model statistics) such that participants who showed greater activation in the PRC to rearranged than intact trials exhibited more behavioural priming. There was no interaction with age, supporting that this was true in both age groups (Fig. 5). That said, only the right PRC model was significant, *F* (3, 50) = 3.83, *p* = 0.015.

## Discussion

Age differences in associative memory are commonly observed when memory is tested explicitly, but not implicitly (Giovanello & Dew, 2015). Despite this difference in performance across test type, the neural correlates of implicit associative memory in aging have largely been ignored. The current study sought to characterize the role of the hippocampus in implicit associative memory and to determine whether hippocampal recruitment during this task differs between younger and older adults. We replicated previous behavioural results showing that older and younger adults did not differ in associative priming (Dew & Giovanello, 2010; Kan et al., 2011), despite differences in next-day explicit associative memory for the same stimuli (Old & Naveh-Benjamin, 2008). Both age groups showed neural associative priming (rearranged > intact) in the left inferior temporal lobe (ITL), left precentral and inferior frontal gyrus (IFG), and right middle frontal gyrus (MFG), in line with the item priming literature (Kim, 2019; Lee et al., 2020). There was also evidence for greater activation in the right middle and inferior temporal lobe and the right and left cuneus for intact compared to rearranged trials in both age groups. Critically, ROI analyses using subject-specific masks of MTL structures indicated that the hippocampus was sensitive to the implicit associative task, but activation patterns differed between age groups. Older adults showed greater activation for intact compared to rearranged trials in the right posterior hippocampus, while young adults showed the reverse pattern in this same region (albeit, not significantly in young; c.f. Gomes et al., 2016; Wang & Giovanello, 2016). While none of the hippocampal ROI contrasts related to behavioural priming (replicating Gomes et al., 2016), greater neural priming in the left and right perirhinal cortex was positively related to greater behavioural priming. Together, these data support past work that implicates the hippocampus in implicit associative memory and further suggests that the perirhinal cortex is also recruited during associative tasks.

Both age groups showed evidence of neural associative priming (rearranged > intact) in the left ITL, left IFG, and right MFG. Each of these regions has been implicated in past research on item priming (Kim, 2019; Lee et al., 2020). The left ITL has been implicated in both the perceptual processing of visual stimuli and more abstract, conceptual representations of those stimuli (e.g., Buckner, 2000; Koutstaal et al., 2001). Our task also included words, thus less activation in the left IFG to repeated pairs might reflect repetition suppression related to processing repeated object words (e.g., Orfanidou et al., 2006; Raposo et al., 2006). However, other work has also implicated this region in response selection and inhibition (Mostofsky & Simmonds, 2008). Given that activation in this cluster also spanned the precentral gyrus, this activation might reflect response selection and inhibition related to the stimulus-response bindings in the object categorization task itself (Horner & Henson, 2009, 2012). However, unlike in item priming tasks, rearranged items were previously paired with other items that might be spontaneously retrieved at test and need to be inhibited. Thus, greater activation in the left IFG, and right MFG (e.g., Nelson et al., 2009), for rearranged than intact pairs may reflect interference resolution (Davis et al., 2024, 2025). Future work should seek to untangle the role of these frontal regions in implicit associative memory tasks and determine how they might differ from item priming paradigms.

The role of the hippocampus in implicit associative memory had been debated (Dew & Cabeza, 2011; Hannula & Greene, 2012; Henke & Ruch, 2024; Steinkrauss & Slotnick, 2024), but arguably, recent work tends to agree that the hippocampus should be involved in tasks that require associative binding regardless of conscious awareness (Cowell et al., 2019). In support of this, Gomes et al. (2016) tested associative priming in young adults using a speeded object categorization task and found greater bilateral posterior hippocampal activation for rearranged than intact pairs at retrieval. Wang & Giovanello (2016) also found task-related activation in the posterior hippocampus in both young and older adults, but their results were reversed (i.e., intact > rearranged). These different effects across the two studies are interesting in the context of our ROI results. Our analysis showed that older adults’ posterior hippocampal activation was greater for intact than rearranged pairs (in line with Wang & Giovanello), while our young adults showed the opposite (in line with Gomes et al.). That said, the young adults’ hippocampal activation in this sample did not differ from zero, nor did their behavioural associative priming scores when analyzed separately (young *p* = .276, old *p* < .001; similar to Davis et al., 2021). While the lack of behavioural evidence in these studies could reflect floor effects in young adults, who respond faster than older adults, the lack of hippocampal effect suggests that some young adults may not have engaged with the object categorization task with the same motivation as older adults (A. D. Ryan & Campbell, 2021).

Nonetheless, it is interesting that our results do not align more closely with Gomes et al., given that our tasks are more similar than with Wang & Giovanello. However, the limited literature on this topic poses a problem in discerning whether we should expect reductions in HPC activation reflecting reduced binding demands to repeated pairs, like in item priming studies (Grill-Spector et al., 2006), or increased HPC activation reflecting the retrieval or reinstatement of previously encoded pairs like in explicit memory studies (Giovanello et al., 2009; Prince et al., 2005). Critically, the consistent posterior hippocampal activation across these experiments indicates that the hippocampus supports implicit associative memory. Future work on this topic should aim to better elucidate the role of the hippocampus in implicit associative memory tasks in aging.

An interesting finding in our study was the relationship between behavioural associative priming and neural associative priming in the perirhinal cortex. The perirhinal cortex is often implicated in studies of item memory likely because of its role in object representations (Clarke & Tyler, 2014; S. Davis et al., 2021), which is a process that is largely preserved with age (Daselaar et al., 2005). Past research has found that the PRC was specifically involved in item, but not associative priming (Gomes et al., 2016; Wang & Giovanello, 2016). That said, there is research in both human and non-human samples to support that the perirhinal cortex may also play a role in associative binding (Haskins et al., 2008; Watson et al., 2012; Yanike et al., 2009), especially in cases where items in a pair can be unitized (Diana et al., 2007; Fiorilli et al., 2021; Giovanello et al., 2006; Haskins et al., 2008;.but see Staresina & Davachi, 2010). The picture-word pairs in this task were not semantically related, unlike in studies that encourage unitization (e.g., word pairs like MOUNTAIN – CLIMBER), but it is possible overlapping picture-word pairs created a situation in which the stimuli could be more easily unitized than if they were spatially separated. However, other work suggests that spatial proximity may not matter (Dennis et al., 2025; but see Seo et al., 2026 [preprint]). Another explanation may be that overlapping stimuli increased demands on visual discrimination, which has also been shown to depend on the PRC (Graham et al., 2010). Though, if this was the case, then we may have expected to see age differences in this PRC process (Ladyka-Wojcik et al., 2026; L. Ryan et al., 2012), which we do not. Thus, while it is possible that the PRC activation evident in our task is related to unitization or visual discrimination, these data also align with studies that implicate the PRC in associative processes (Naya, 2016). Future work should aim to determine the context in which PRC activation supports implicit associative memory and when it does not.

These data fit with a larger literature regarding the role of the MTL in memory in aging. Preserved performance on implicit associative tasks have been used to argue that older adults’ deficits in explicit associative tasks are a result of frontally-mediated control processes rather than associative binding (Campbell & Davis, 2024; Giovanello & Dew, 2015; Hasher et al., 2007). Indeed, several studies support that age-related changes to prefrontal structure and function affect explicit associative memory in aging (Addis et al., 2014; de Chastelaine et al., 2016; Guardia et al., 2023). Here, we support these theories by demonstrating that if we minimize the need for these frontally-mediated control processes by testing memory implicitly, then there is both behavioural and neural evidence that older adults form associations between two unrelated stimuli. While the implicit retrieval of these associations appears to be supported by the hippocampus, its role still may differ in young and old adults. Given how few studies have examined this question, replicating these findings is essential for understanding implicit associative memory and how it differs from explicit memory in aging. Identifying why implicit and explicit memory diverge with age may, in turn, point to ways of supporting older adults’ performance on explicit memory tasks.

## Author Contributions

Designed research (E.E.D, S.W.D, K.L.C.), performed research (E.E.D., K.L.C), analyzed data (E.E.D), contributed analytic tools (R.K.O), wrote the first draft of the paper (E.E.D), wrote the paper (E.E.D., K.L.C), edited the paper (E.E.D, S.W.D, R.K.O., K.L.C).

## Conflicts of Interest

The authors report no conflicts of interest.

## Acknowledgments

We would like to thank research assistants H.T., L.A., and I.C. for their help with data collection. This work was supported by Natural Sciences and Engineering Research Council of Canada (NSERC) Postdoctoral Fellowship award (E.E.D), and NSERC Discovery Grant (K.L.C).

## Notes

### Competing Interest Statement

The authors have declared no competing interest.

