## Supplemental Table 1 for "Not all memory declines with age: Neural correlates of implicit associative memory in younger and older adults"

**Supplemental Table 1. Regression models predicting behavioural associative priming from age group and neural associative priming (rearranged > intact) in each MTL ROI.**

| | **ROI slope** | | | **Age group** | | | **ROI x Age** | | | **Model** | | |
|---|---|---|---|---|---|---|---|---|---|---|---|---|
| **Region** | *b* | *SE* | *p* | *b* | *SE* | *p* | *b* | *SE* | *p* | $R^2$ | *F* | *p* |
| L anterior HPC | 25.13 | 14.50 | 0.089 | 20.49 | 16.50 | 0.220 | -28.12 | 29.00 | 0.337 | 0.100 | 1.85 | 0.150 |
| R anterior HPC | 16.51 | 14.99 | 0.276 | 23.33 | 16.79 | 0.171 | -21.49 | 29.98 | 0.477 | 0.069 | 1.23 | 0.307 |
| L posterior HPC | 1.53 | 23.12 | 0.947 | 20.90 | 17.02 | 0.225 | -43.12 | 46.24 | 0.356 | 0.046 | 0.80 | 0.499 |
| R posterior HPC | 23.66 | 24.67 | 0.342 | 26.16 | 18.12 | 0.155 | -36.54 | 49.33 | 0.462 | 0.052 | 0.91 | 0.441 |
| L PHC | 14.16 | 16.57 | 0.397 | 20.64 | 16.95 | 0.229 | 21.42 | 33.13 | 0.521 | 0.050 | 0.88 | 0.460 |
| R PHC | 20.66 | 16.87 | 0.226 | 23.00 | 16.74 | 0.176 | 18.33 | 33.73 | 0.589 | 0.071 | 1.28 | 0.293 |
| L PRC | 46.95 | 19.33 | **0.019*** | 19.68 | 16.22 | 0.231 | -6.73 | 38.66 | 0.863 | 0.132 | 2.54 | 0.067 |
| R PRC | 50.23 | 16.58 | **0.004*** | 27.03 | 15.74 | 0.092 | -32.96 | 33.16 | 0.325 | 0.187 | 3.83 | **0.015*** |

*Note.* HPC = Hippocampus. PHC = Parahippocampal cortex. PRC = Perirhinal cortex.

*$p < .05$
